# Beauty at risk: A taxonomic synopsis of *Belemia* (Nyctaginaceae), an endangered and endemic genus of vines in Brazil

**DOI:** 10.64898/2026.05.12.724086

**Authors:** Israel L. Cunha-Neto, Elson Felipe S. Rossetto, Daniel V. Goncalves, Matheus G. C. Nogueira, Guilherme M. Antar, Victor R. C. Rodrigues, Alessandro O. Silva, Veronica Angyalossy, Cyl Farney C. Sá

## Abstract

*Belemia* belongs to Nyctaginaceae and comprises two species of delicate vines. Both species are endemic to Brazil. *Belemia fucsioides*, the type species, described in 1981, occurs in a restricted area of the Atlantic Forest in southeastern Brazil. *Belemia cordata*, described in 2020, is known from only two records from the same area in the Cerrado of northern Brazil. Here, we describe the taxonomic history of *Belemia* and provide the first synopsis for the genus. We include species description, distribution map, identification key, and anatomical data. We used field observations over the past decade and modeled anthropogenic changes in the species range to conduct a conservation assessment in accordance with the IUCN Red List criteria. Conservation assessments indicate significant concerns for *Belemia*, classified as either endangered (*B. fucsioides)* or critically endangered *(B. cordata)*. The species are threatened primarily by habitat loss to land used for agriculture, forestry, and livestock production. This study contributes to ongoing initiatives exploring plant diversity in the Neotropics and supports efforts to identify threats to biodiversity.

## Introduction

*Belemia* Pires is a small genus in Nyctaginaceae, endemic to Brazil, and characterized by delicate vines (climbing plants) bearing tubular, colorful flowers (Flora e Funga do Brasil 2026). Molecular data placed this genus as a sister lineage to *Phaeoptilum* Heimerl, a monotypic genus endemic to Africa, which emerges as sister to the Neotropical and widely cultivated *Bougainvillea* Comm. ex Juss. Together, these three genera form the tribe Bougainvilleae (Douglas and Spellenberg 2010). Despite clarification of their phylogenetic relationship, a morphological synapomorphy for the tribe Bougainvilleeae has not been proposed so far.

*Belemia* was the last genus of Nyctaginaceae to be described, with the type species, *B. fucsioides* Pires, being described from southeastern Brazil in 1981 (Pires 1981), and the second, *B. cordata* Harley & Giul., described in 2020 from a single locality in the Cerrado of northern Brazil (Zappi *et al*. 2020). Vegetatively, *Belemia* differs from *Bougainvillea* and *Phaeoptilum* primarily because of alternate leaves and a perianth constricted above the ovary, as well as the absence of spines (Bittrich and Kubitzki 1993). The scarce records of *Belemia* species in herbaria and online databases have made it little known within the scientific community, and this paper aims to address this gap and foster opportunities for the conservation of these remarkable climbing plants.

Here, we provide an overview of *Belemia*, summarizing its history and taxonomy. We include species distribution maps, images from field collections, an identification key, and anatomical data. Additionally, we evaluate each species’ distribution and assess its conservation status.

## Material and Methods

### Taxonomic study

The study was based on a literature review, herbarium studies, and field collection efforts, primarily in areas with a high chance of finding *Belemia* species based on previous field expeditions. Field expeditions were conducted in 2008, 2018, 2021, and 2026 in the occurrence area of *B. fucsioides*, and in 2016 and 2026 in the area of *B. cordata* (Supplementary Table S1). In the field, observations and photographs were taken of each species during the vegetative, flowering, and fruiting periods. Fresh samples were collected for anatomical studies during the 2021 field expeditions. In addition to new collections, we examined specimens deposited at the RB, SAMES, and NYBG herbaria (Index Herbariorum (updated continuously) 2026). Additional specimens deposited in other herbaria were analyzed using vouchers from online repositories, including *SpeciesLink* (SpeciesLink 2026) (https://specieslink.net/), REFLORA (Flora e Funga do Brasil 2026) (https://floradobrasil.jbrj.gov.br/reflora), JABOT (JBRJ - Instituto de Pesquisas Jardim Botânico do Rio de Janeiro 2026) (https://rb.jbrj.gov.br/v3/consulta.php), or GBIF (GBIF.org 2026) (https://www.gbif.org/). Species descriptions and an identification key were elaborated primarily based on morphological comparisons among species indicated in Supplementary Table S2.

### Conservation assessment

Distribution maps for each species were prepared in QGIS LTR 3.40 Software (QGIS.org. 2026) using geographic coordinates collected in the field for each individual, and shapefiles for Brazil’s Biomes and Federal Units available at Instituto Brasileiro de Geografia e Estatística (IBGE 2026a; b). Land-use maps were created using satellite imagery (Copernicus Sentinel data 2026). The Sentinel data was modified in QGIS using the Semi-Automatic Classification Plugin (Congedo 2021). The conservation assessment of each species is according to the IUCN Red List categories and criteria (IUCN 2024), and the extent of occurrence (EOO) and area of occupancy (AOO) values were calculated using the GeoCAT tool (https://geocat.iucnredlist.org) (Bachman *et al*. 2011) using a default cell area of 2 km^2^. To estimate the EOO and AOO for *B. cordata* (which has only two geographical coordinates), we included a third coordinate near the location where the species was collected in 2026.

### Anatomical study

During the 2021 field expedition, we collected fresh, fully expanded leaves and stems from different developmental stages of *B. fucsioides* (*I. L. Cunha-Neto 17*) for anatomical studies. Anatomical studies were also conducted using a leaf and stem sample from a voucher of *B. cordata* (*M. G. C. Nogueira 544*). Leaf and stem samples were processed using standard plant microtechnique protocols (Angyalossy *et al*. 2016), either by hand sectioning (for small samples) or using microtome sectioning (for larger stem samples). Microscope slides were imaged using a Leica DMBL light microscope coupled with a digital camera (Leica DFC310, Leica Microsystems, Wetzlar, Germany).

## Results

### History of *Belemia*

Although their showy flowers are easily spotted in the field (Fig. 1), there have been only a few unique records of *Belemia* in herbaria, and most of the 23 records in SpeciesLink (SpeciesLink 2026) and 36 in GBIF (GBIF.org 2026) are duplicates distributed from only 15 herbarium collections (Supplementary Table S1). Of the 15 collections, two were of *B. cordata* and 13 of *B. fucsioides*; these observations were conducted by 12 different collectors (Supplementary Table S1). *Belemia* has distinctive features that facilitate its field observation and taxonomic description; thus, the low number of records is likely due to its narrow distribution and habitat loss within its range.

**Figure 1.**
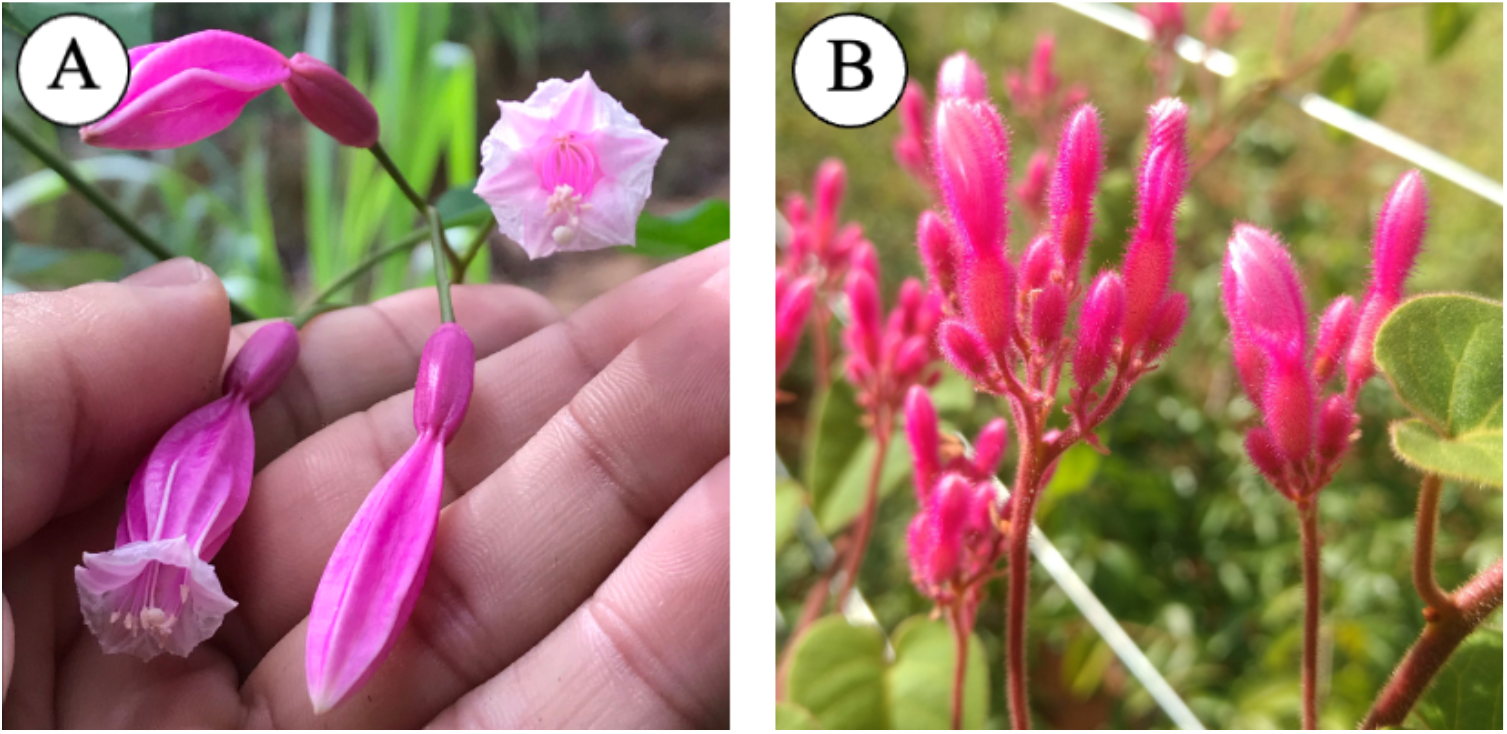
The showy flowers (and flower buds) of *Belemia* species. (A) *Belemia fucsioides*. (B) *Belemia cordata*.

The formal description of the type species (*B. fucsioides*) occurred in 1981 (Fig. 2); however, the oldest herbarium specimen of this species dates to 1953 (*A. P. Duarte 3704*), and this and other herbarium specimens remained undetermined for years in Brazilian and foreign herbaria. One of these collections (*J. P. Lanna Sobrinho 1003*) was made near the type locality in the far northern part of Espírito Santo state (southeastern Brazil), the day before the holotype was collected in 1965. Another collection (*R. P. Belém 3796)* occurred in 1968 between the municipalities of Nanuque and Teófilo Otoni, state of Minas Gerais (MG), prior to the description of the genus.

**Figure 2.**
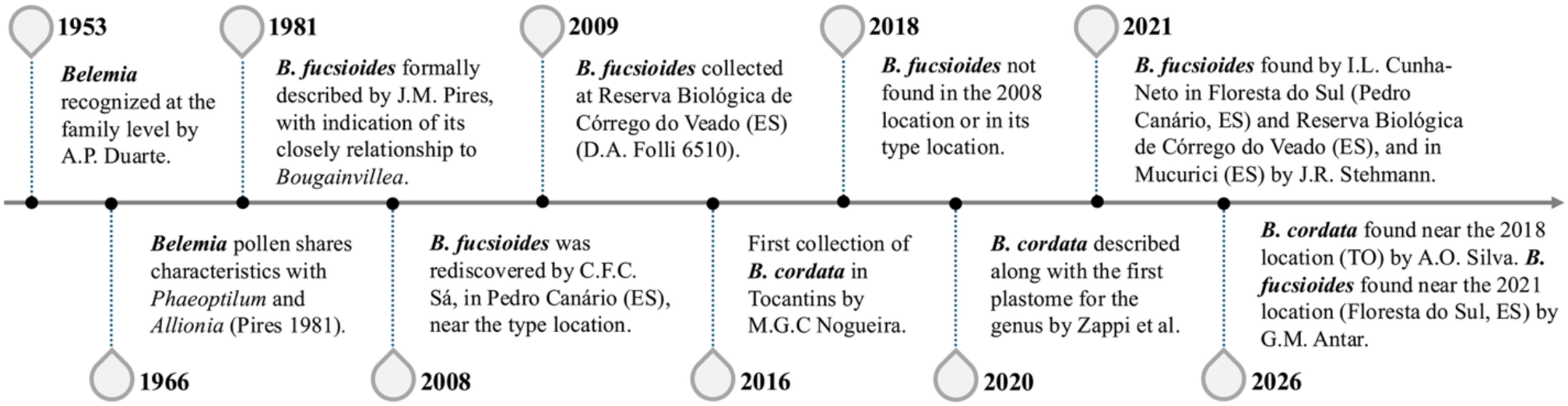
Timeline of some of the important events in the taxonomic history of *Belemia*. (Inspired by Zeferino et al., 2025; Acta Bot. Bras.).

After a few decades since its description, *B. fucsioides* was collected again (*C. F. C. Sá 4887, 4888*) in the municipality of Pedro Canário, Espírito Santo state, near a forest fragment surrounding a sugarcane plantation along the BR 101 highway, during an expedition to locate the species in 2008 (Figs. 2, 3). The following year, one specimen (*Folli 6510*) was collected in the Reserva Biológica de Córrego do Veado, Pinheiros, Espírito Santo, approximately 30 km from Pedro Canário. Nearly ten years later (2018), a new attempt to collect the species for anatomical studies failed to locate the plant at the same site in Pedro Canário, as the sugarcane plantation area had expanded into the forest fragment where it had been collected in 2008.

**Figure 3.**
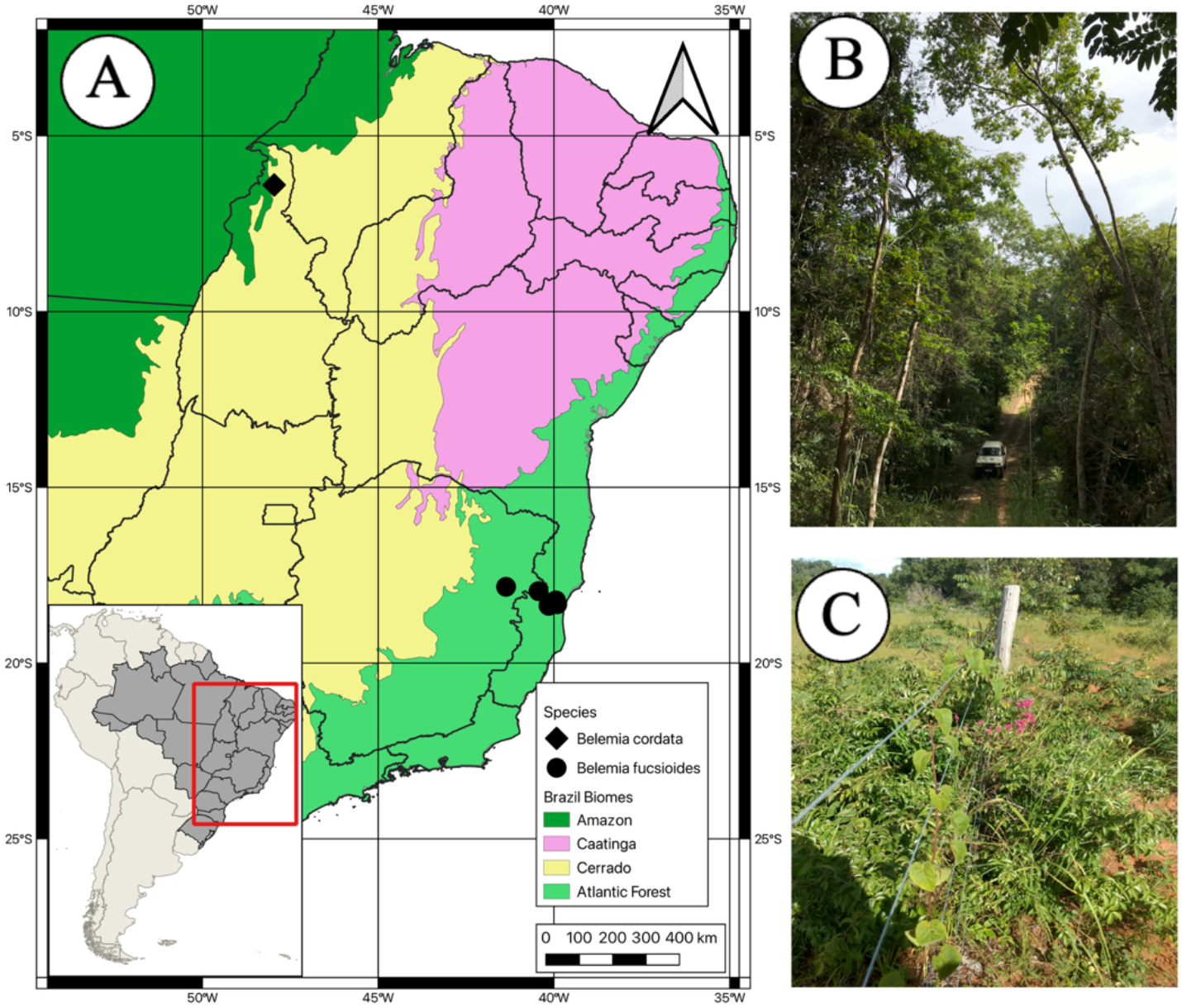
Distribution and habitat of *Belemia*. (A) Map of the two *Belemia* species and their corresponding biomes. (B) Atlantic Forest in Espirito Santo (Reserva Biológica Córrego do Veado), where *Belemia fucsioides* are found. (C) Cerrado in Tocantins, where *Belemia cordata* are found.

In 2021, another expedition to collect the species recorded two observations in Espírito Santo: one in the Reserva Biológica de Córrego do Veado (*I. L. Cunha-Neto 16*), in the municipality of Pinheiros, and another near the village of Floresta do Sul towards the valley of the Rio Itaúnas (*I. L. Cunha-Neto 17*), situated between Pinheiros and Pedro Canário. That same year, *B. fucsioides* was collected about 80 km away, in the municipality of Mucurici, Espírito Santo (*J. R. Stehmann 6568*). Prior to the submission of this manuscript, we attempted to observe *B. fucsioides* in the two areas near Pedro Canário corresponding to the specimens collected in 2008 and 2021 (Floresta do Sul) and found it only in the latter (*G. M. Antar 6212*). The scarce information on *B. fucsioides* led to its categorization as “Endangered (EN)” in the List of Endangered Fauna and Flora – Espírito Santo (Fraga *et al*. 2019).

*Belemia cordata*, the second species described in the genus, was first recorded in 2016, and its description was published a few years later in a study that also provided the first plastome for the genus (Zappi *et al*. 2020). The 2016 collection of *B. cordata* was made during fieldwork along the TO-210 road between Ananás and Angico, where the plant was first noticed because of its conspicuous pink flowers growing in disturbed Cerrado vegetation. This area is a highly fragmented Cerrado known for agrarian conflicts in the state of Tocantins, Brazil (Fig. 3). The species was described based on specimens from that single location (Zappi *et al*. 2020). *Belemia cordata* is distinguished from *B. fucsioides* primarily by leaf shape, flower color, and shoot pubescence (Zappi *et al*. 2020). Nevertheless, they share a tragic condition of being endangered, even though they are found more than 1,000 km apart in different biomes (Zappi *et al*. 2020) (Fig. 3). There had been no reports of observations of this plant since its publication, but our efforts to observe it in the field before submitting this manuscript resulted in a new collection at the type locality (*Silva 130*), despite ongoing human activities in the area.

### Taxonomic treatment

***Belemia*** Pires, Bol. Mus. Paraense Emílio Goeldi, N.S., Botânica 52: 1. 1981.

#### Type

*Belemia fucsioides* Pires

Lianas, glabrescent (*B. fucsioides*) to puberulous (*B. cordata*), including glandular trichomes; tubercles present in *B. fucsioides* and not found in *B. cordata*. Branches terete, fistulose.

Leaves simple, alternate, being larger at base and decreasing in size toward the apex of branches; petioles twining; blades ovoid with rounded to asymmetrical base (*B. fucsioides*) or deltoid with cordate base (*B. cordata*), apex acuminate, midrib pinnate, impressed on the adaxial surface and prominent on the abaxial surface, secondary venation brochidodromous, impressed on the adaxial surface and prominent on the abaxial surface. Inflorescences axillary and terminal, usually dichotomously branched, occasionally unbranched, with branch scars and bracts along its length, each primary branch terminating in a solitary, sessile flower. Flowers magenta or pink in the field, bisexual, with a constriction in the middle portion of the perianth, being cylindrical, thickened below the constriction, membranaceous, 5-6 lobed above; stamens 10-12, basally conate, reaching the perianth opening or slightly exserted, anthers basifix, longitudinal slits; stigma included, subglobose, and papillose.

Anthocarps elliptic to narrowly ovoid, coriaceous, cylindrical, filaments persistent, portion above the constriction curling up.

#### Etymology

The generic epithet honors Romeu Pedro Belém, one of the first field assistants of the CEPEC Herbarium (1965–1980) and the collector of the type of *Belemia* (Mori and Silva 1980).

#### Notes

The distinctive characters that distinguish the genus from *Bougainvillea* and *Phaeoptilum* include spineless, slender vines with twinning petioles; showy pink to magenta flowers without an involucrum; and a perianth constricted in the middle.

### Key to the species of *Belemia*

1. Plants glabrescent, inconspicuous glandular trichomes, leaf base rounded or asymmetrical, bracts of inflorescences minute. Atlantic Forest of northern Espírito Santo and northeastern Minas Gerais states................ *Belemia fucsioides*

1’. Plants puberulous, glandular trichomes visible, leaf base cordate, bracts of inflorescences showy and foliaceous. Cerrado of Northern Tocantins state..................... *Belemia cordata*

1. ***Belemia cordata*** Harley & Giul., Syst. Biodiv. 18(4): 331. 2020. (Figs. 4, 5). Type: BRAZIL. Tocantins: between Angico and Ananás, along the TO-210 road, 9 July 2016, *M*.*G*.*C. Nogueira et al. 544* (holotype: MG; isotypes NY, HUEFS, RB).

**Figure 4.**
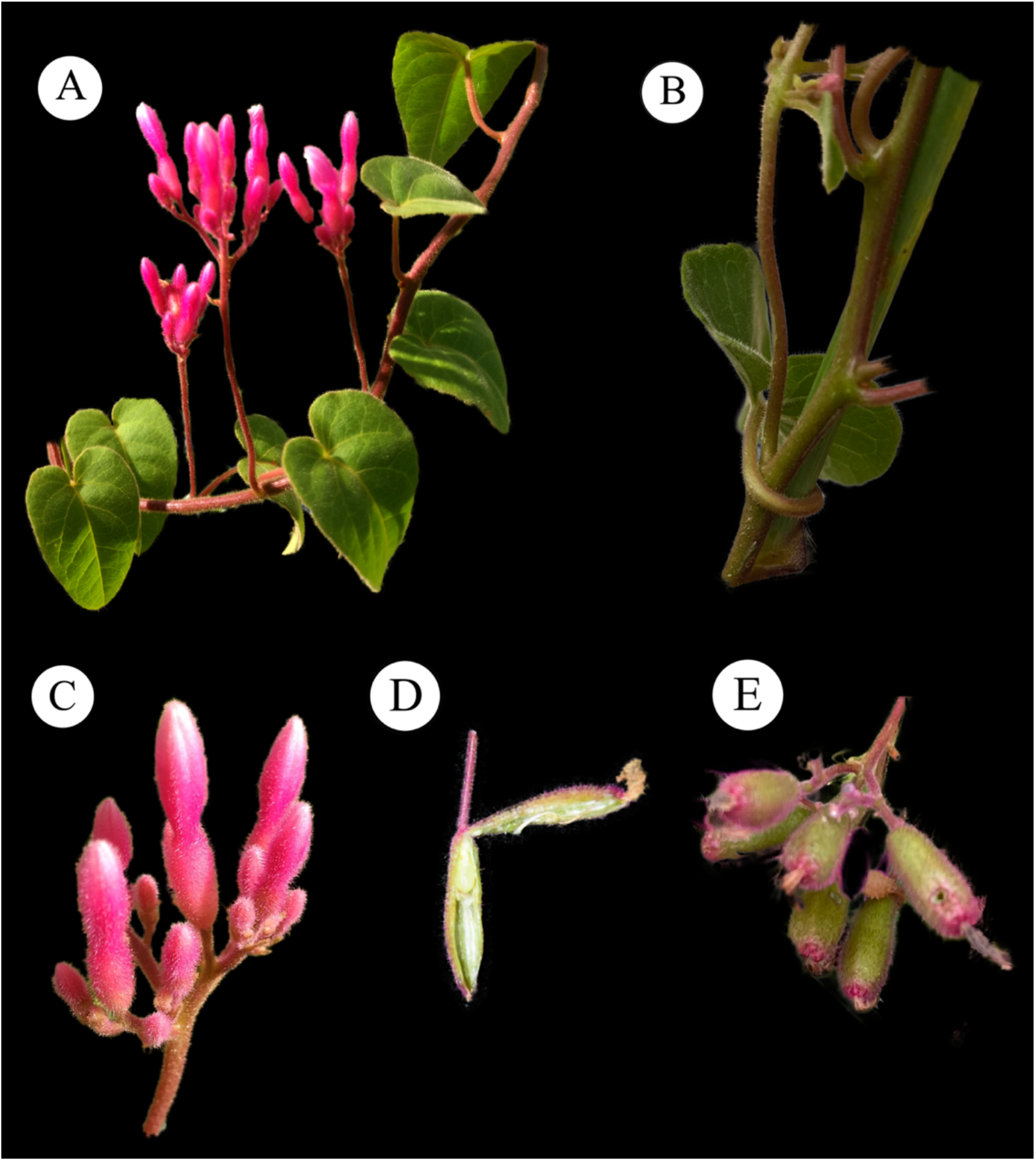
Morphological characters of *Belemia cordata*. (A) Climbing habit. Cordate leaves. Sub-terminal axillary inflorescences. (B) Twinning petiole (climbing mechanism). (C) Light pink flower buds. (D) Dissected flower. (E) Anthocarp.

**Figure 5.**
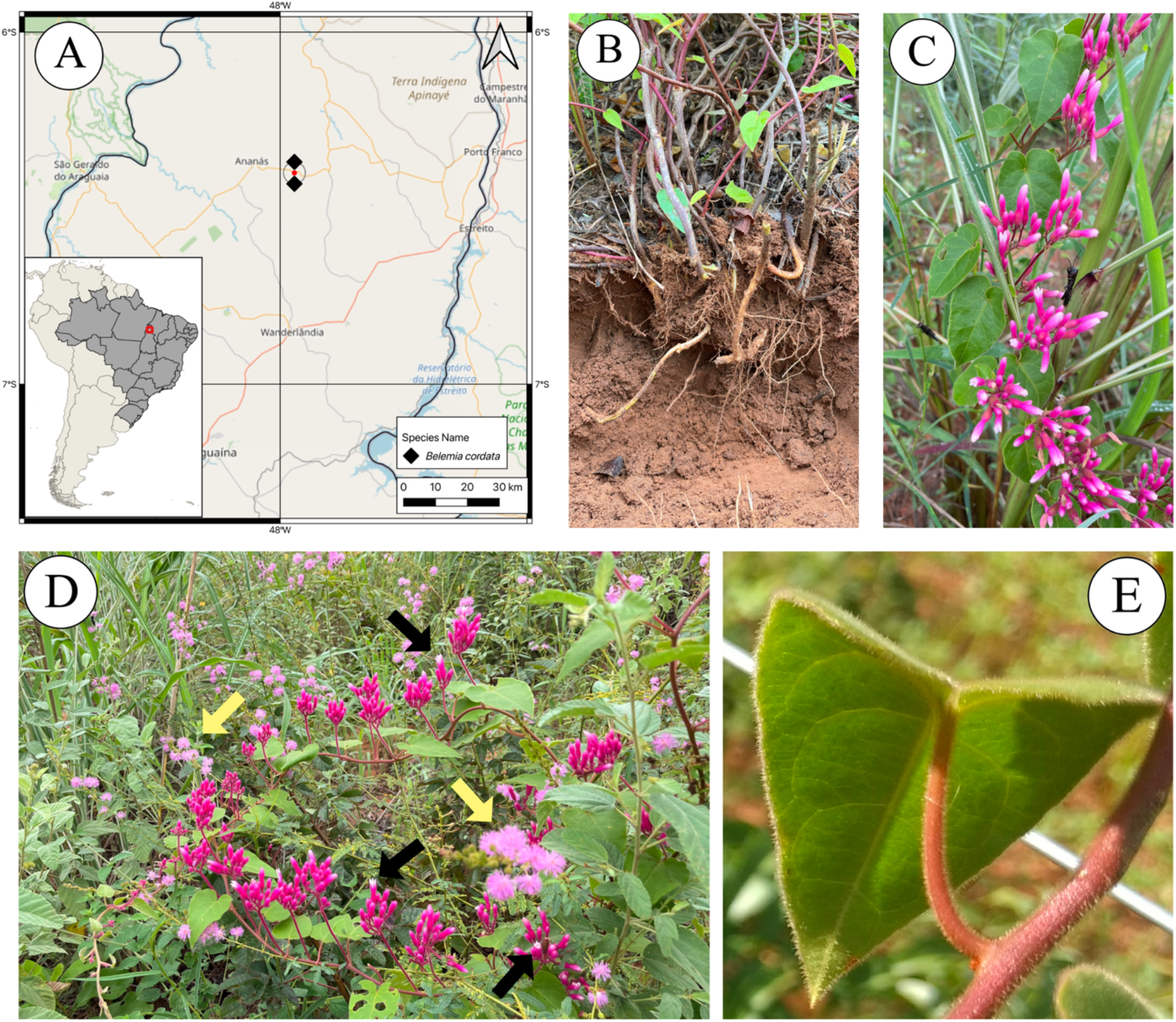
Distribution map and field observations of *Belemia cordata*. (A) Distribution map. (B) Base of the plant showing a lack of a subterranean organ. (C) Cordate leaves and inflorescence. (D) Climbing habit. Note open flowers from *B. cordata* (black arrow) and from a Fabaceae species (yellow arrow). (E) Young cordate leaf, with non-twining petiole.

#### Description

Twining petiole climbers, mostly puberulous. Tubercle not found. Leaf blades 1.0-13.0 × 1.0-7.0 cm, deltoid, base cordate, apex acuminate, chartaceous, concolor, margin ciliate; secondary veins 6-8 pairs, diverging from the primary vein at an angle of 50º in the middle of the blade and 90º at the base, intersecondary veins 1-2, tertiary veins conspicuous in the lower surface. Inflorescence erect; peduncle 0.3-6.0 cm long, primary branches 0.1-1.9 cm long; bracts showy and foliaceous. Flowers pink in the field, 0.8-2.6 cm long; filaments 0.8-1.2 cm long, anthers 1-1.2 mm long; style 0.8-1.2 cm long. Anthocarps 1.5-1.9 × 0.3-0.5 cm.

#### Etymology

The specific epithet refers to the outstandingly cordate leaves (Zappi *et al*. 2020).

#### Distribution and habitat

*Belemia cordata* was recorded exclusively in the state of Tocantins, Brazil, with occurrences along the TO-210 highway, near the municipality of Angico and Ananás. The area lies within the Cerrado, at the ecotone between the Cerrado and the Amazon biomes.

#### Phenological note

Observed with flowers in July and March. Fruits were observed in March.

#### Specimens examined

BRAZIL. **Tocantins:** margem da rodovia TO-210, entre Angico e Ananás, 27 Feb 2026, bd., fl., fr., *A*.*O. Silva 130* (IFTO)

#### Conservation status

Critically Endangered (CR). *Belemia cordata* is known only from two collections in the same area in the state of Tocantins, Brazil. In the 2026 field expedition, only one voucher specimen (*Silva 130*) was produced, yet more than one individual was observed in the same area. Its conservation status is considered Critically Endangered (CR B2ab(ii, iii)). Since the EOO is smaller than the AOO, they are estimated at the same value: 8 km^2^. Inference about the potential distribution of this species is difficult due to many unknowns regarding its biology and habitat requirements. This species faces a variety of threats, but the most pressing is the conversion of its habitat to agricultural land. This makes the likelihood of this population surviving very low.

2 ***Belemia fucsioides*** Pires, Bol. Mus. Paraense Emílio Goeldi, N.S., Botânica 52: 2. 1981. (Figs. 6, 7).

**Figure 6.**
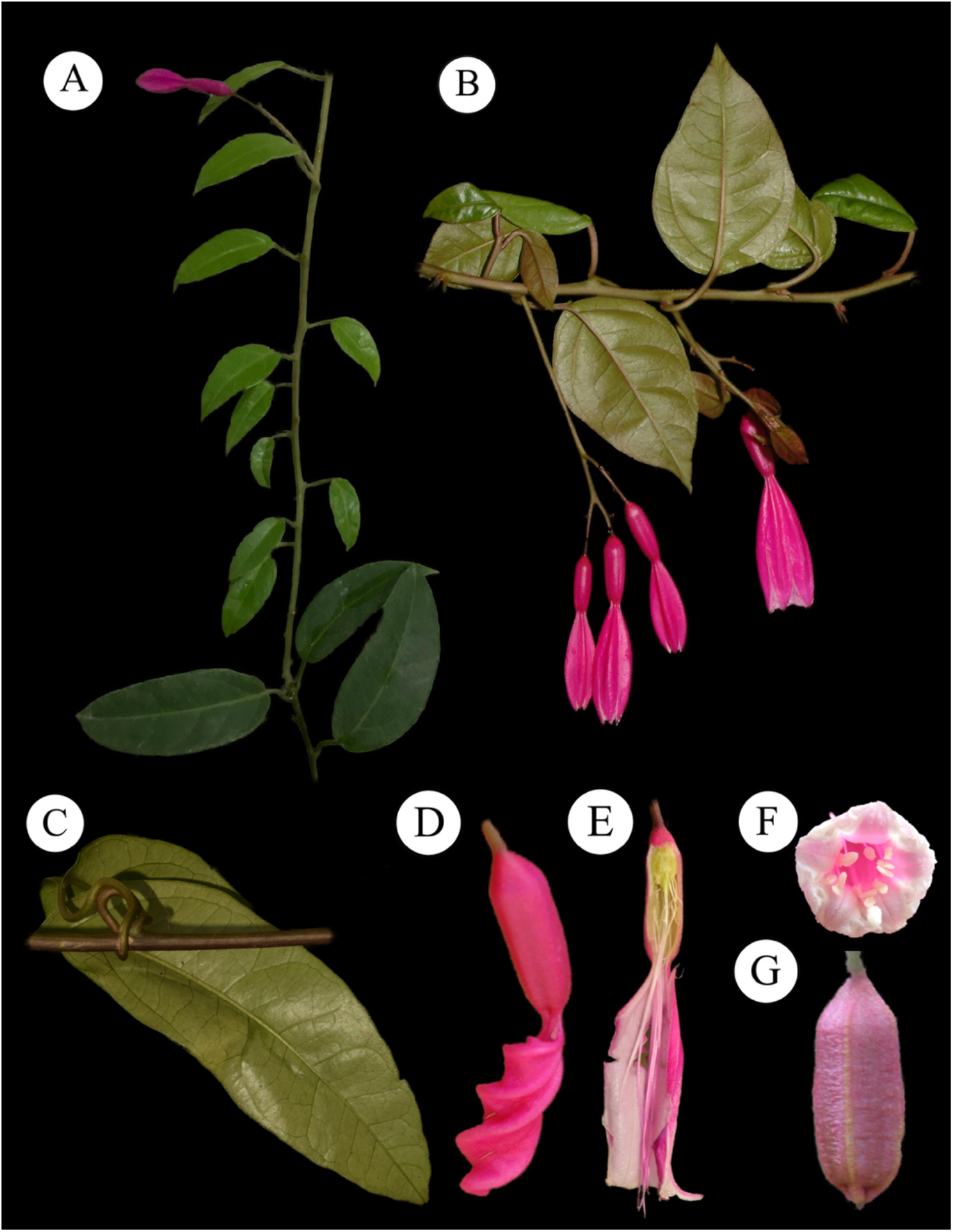
Morphological characters of *Belemia fucsioides*. (A) Branch with leaves and inflorescence. (B) Lanceolate leaves and flowers with a constricted region. (C) Twinning petiole. (D) Early anthocarp development with twisted, persistent perianth. (E) Open flower showing stigma and stamens. (F) Frontal view of flowers. (G) Anthocarp.

**Figure 7.**
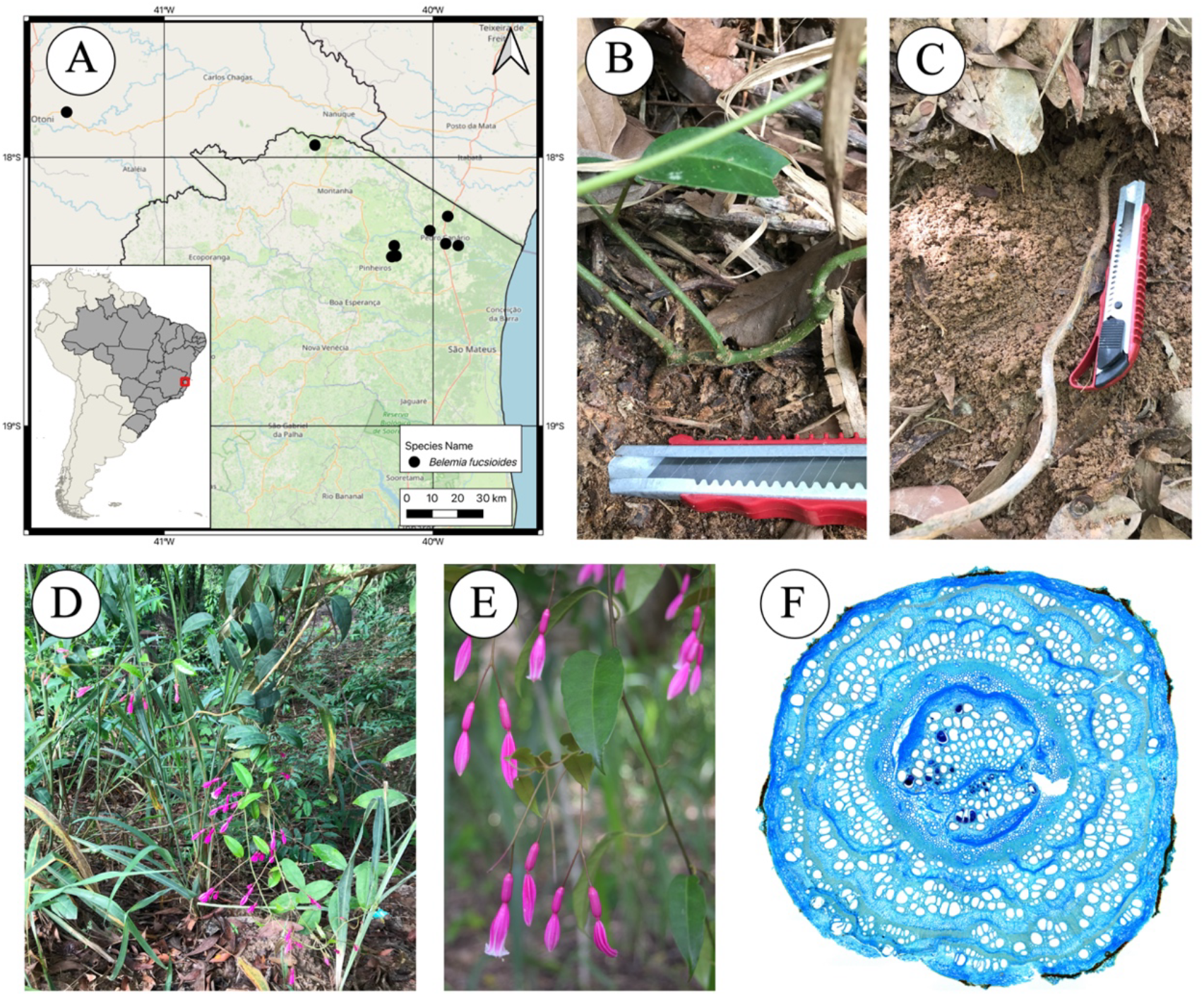
Distribution map and morpho-anatomical observations of *Belemia fucsioides*. (A) Distribution map. (B) Base of the stem aboveground. (C) The base of the stem underground. (D) Climbing habit. (E) Flowers. (F) Cross-section of an adult stem; stem diameter = 10 mm.

Type: BRAZIL. Espírito Santo: [Pedro Canário], BR-5 [currently BR-101], 5 km ao sul do Morro Dantas, beira da estrada, 9 Aug 1965, *R*.*P. Belém 1460* (holotype: MG barcode MG071318_1 [image!]; isotypes: NY barcode NY00342017 [image!], UB barcode UB00040107 [image!]), CEPEC 1289, IAN 119786.

#### Description

Twining petiole climbers, glabrescent, with a higher concentration of inconspicuous capitate-stalked trichomes on bracts, buds, and the branching points of branches. Tubercle present. Leaf blades (0.3-) 1.7-14.5 × (0.4-) 2.0-6.5 cm, ovate, base rounded or asymmetrical, apex acuminate, chartaceous, concolor, margin glabrous, secondary veins in 3-7 pairs, diverging from the primary vein at an angle of 40º-70º in the middle of the blade and 45º-50º at the base, intersecondary veins 1, tertiary venation conspicuous in lower surface. Inflorescences pendulous; peduncle 1.5-9.0 cm long, primary branches 0.1-1.3 cm long; bracts diminute. Flowers magenta in the field, 3.1-4.5 cm long; filaments 3.2-4.7 cm long, anthers 1.3-1.8 mm long; style 3.2-4.5 cm long. Anthocarps 1.5-1.8 × 0.4-0.6 cm.

#### Vernacular name

cipó-sino (*D. A. Folli 6510*).

#### Etymology

The specific epithet likely refers to the form and color of *Fuchsia* (Onagraceae) flowers. However, no reference to that was proposed in the original description (Pires 1981). **Distribution and habitat:** *Belemia fucsioides* has been recorded only in Tabuleiro Forests of the northernmost portion of Espírito Santo state, specifically in the municipalities of Mucurici, Pedro Canário, and Pinheiros. This vegetation formation, also known as the Coastal Tablelands (“Tabuleiros Costeiros”), is part of the Atlantic Forest biome, being humid but with seasonal influence (Peixoto *et al*. 2008), and covers the Cenozoic sedimentary lowlands along the coast of Northern Espírito Santo and Southern Bahia.

Moreover, there is one occurrence reported in an area of semideciduous seasonally forest of Minas Gerais state (ca. 180 km from the type locality in a straight line) (Figs. 3, 6).

#### Phenological note

Observed with flowers in February, March, June, July, August, October, and November. Observed with fruits in October and December.

#### Specimens examined

BRAZIL. **Espírito Santo**: Mucurici, Distrito de Água Boa, interior de mata, 26 Nov 2021, bd., fl., *J*.*R. Stehmann et al. 6568* (BHCB barcode BHCB207518, RB barcode RB01502864!); Norte do Espírito Santo [Pedro Canário], entre o Córrego da Preguiça e Rio Preto, bd., fl., fr., 6 Nov 1953, *A*.*P. Duarte 3704* (RB barcode RB00265942!); Morro Dantas [currently Pedro Canário], BR-5 [currently BR-101], 8 Aug 1965, fl., fr., *J*.*P. Lanna Sobrinho 1003* (RB barcode 01281517!); Pedro Canário, estradas vicinais próximas ao eixo da BR-101 entre o Rio Itaúnas e 5 km em direção a Pinheiro, 21 Oct 2008, bd, fl., *C. Farney et al. 4887* (RB barcode RB00943854!); same locality, 21 Oct 2008, bd., fl., *C. Farney et al. 4888* (RB barcode 00943856!); same locality, estrada para Floresta do Sul, 28 Feb 2021, bd., *I*.*L. Cunha-Neto 17* (RB barcode RB01464990!, SPF); same locality, 21 Jan 2026, fl., *G*.*M. Antar & L. Fernandez 6212* (VIES). Pinheiros, Reserva Biológica Córrego do Veado. 09 Dec 2009, bd., fl., fr., *D*.*A. Folli 6510* (CVRD, RB barcode 00856866!); same locality, 07 Nov 2022, fl., *P*.*M. Nunes et al. 101* (SAMES). **Minas Gerais**: margem da Rodovia Teófilo Otoni-Nanuque, zona calcárea, 30 Jun 1968, bd., fl., *R*.*P. Belém 3796* (NY barcode NY00642291 [photo!]).

#### Conservation status

Endangered (EN). *Belemia fucsioides* is known to inhabit a small area in southeastern Brazil. Its conservation status is considered Endangered (EN A2c; B1ab(i,ii,iii)+B2ab(i,ii,iii), with an EOO of 2993.64 km^2^, and an AOO of 32 km^2^. Three of the most recent observations of this species were made inside the protected area of Reserva Biológica de Córrego do Veado, and two were found outside. One of the populations in the municipality of Pedro Canário (road to Floresta do Sul) is particularly threatened, as it occurs at the edge of a small forest fragment that is progressively being invaded by *Acacia mangium* Willd., one of the most aggressive invasive species in the region (Lehmann *et al*. 2017).

*Belemia fucsioides* faces a variety of threats, but the most pressing is habitat conversion into agricultural land and habitat fragmentation. This makes crossbreeding between individuals harder and reduces their chances of survival. The only specimen known from Minas Gerais state was collected in 1968, with no records made since then. Although our 2018 expedition failed to rediscover populations in the area, further attempts are necessary to relocate the species or confirm its extinction in this state.

### Anatomical notes

Leaves have uniseriate epidermis, formed by flattened, oval to round cells, covered by evident cuticle; stomata are observed on the abaxial surface, at the same level as the other epidermal cells (Supplementary Figure S1). Glandular trichomes are sparse in *B. fucsioides* (observed only under a stereoscope and in anatomical sections) and abundant in *B. cordata*, densely covering both leaf surfaces. The mesophyll is dorsiventral with one layer of short palisade parenchyma and a compact spongy parenchyma formed by isodiametric cells, and collateral vascular bundles arranged in the middle portion. The midrib and petiole contain lamellar collenchyma and isodiametric parenchymatic cells in the cortex, and the vascular system is formed by three larger bundles facing the abaxial side, and three to four smaller bundles towards the adaxial side in the midrib; the petiole contains 8 to 12 bundles arranged in a ring (Supplementary Figure S1). Prismatic crystals occur in pith cells.

Stems have an uniseriate epidermis and a cortex composed of collenchyma and parenchyma (Supplementary Figure S1). The vascular system exhibits the common variants observed in Nyctaginaceae (Cunha Neto *et al*. 2020, 2022), including “medullary bundles”— additional vascular bundles in the pith of the stem—that form during early developmental stages, and “interxylary phloem”— alternating bands of secondary xylem and secondary phloem derived from the atypical activity of a single vascular cambium—which develops in adult stems (Fig. 7; Supplementary Figure S1). The secondary xylem displays typical vine anatomy, with many large vessels from the start of secondary growth, indicating a clear transition from a previous self-supporting phase with fewer vessels and more fibers (Supplementary Figure S1). Mature stems produced evident lenticels; however, most of the stem circumference retains the epidermis (Supplementary Figure S1).

## Discussion

In this study, we provide a revision of *Belemia*, revealing the diagnostic characters for each of the two species, their occurrence range, habitat, and conservation status. Although our study does not identify traits that could be potential synapomorphies for Bougainvilleae, we note that the twisted, persistent perianth observed in *Belemia* (Fig. 6D) also occurs in *Bougainvillea* (Heimerl 1934). We have not gathered sufficient data (from the literature or public databases) to determine whether this phenomenon also occurs in *Phaeoptilum*.

The conservation assessment of *Belemia* species provided concerning results, as they are critically endangered (*B. cordata*) and endangered (*B. fucsioides*) species, threatened mostly by habitat loss to land dedicated to agriculture, forestry, and livestock (Fig. 8).

**Figure 8.**
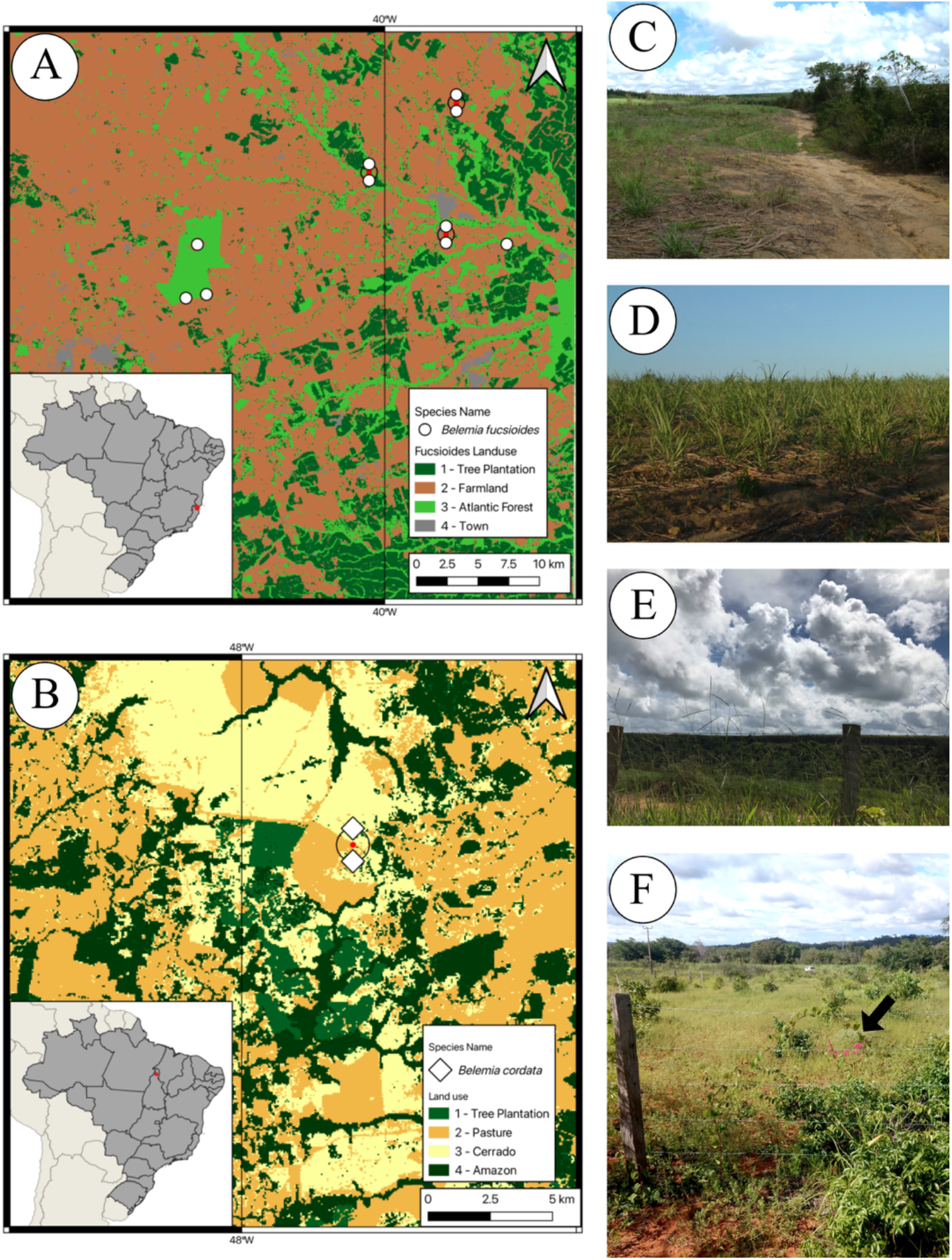
Land-use map for the range of *Belemia* species, and sites where plants were found. (A) *Belemia fucsioides*. (A-B). Land-use maps. The red circle anchors two closely spaced geographic coordinates. (B) *Belemia cordata*. (C-D) Farmland near *B. fucsioides*. Note sugarcane plantation in D. (E) Eucalyptus plantation (in the background) near *B. fucsioides*. (F) Pasture near *B. cordata* (arrow).

Initially, we hypothesized that *Belemia* could have a broader distribution, including more populations escaping areas affected by the growing anthropogenic pressures. We predicted that specimens might have been misfiled in the wrong cabinets, mixed with other common genera. Indeed, the genus has a history of specimens being misfiled into other families with showy tubular flowers, including the type specimen, which was placed under Passifloraceae (*R. P. Belém 1460*), and subsequent collections that were found under Acanthaceae (*D. Folli 6510*) or Convolvulaceae (*P. M. Nunes 101*).

Despite the identification of misfiled herbarium specimens, the recent collections did not expand the group’s known distribution and simply confirmed records within its established range. The limited distribution range of *Belemia* may be due to the absence of accessory or anthocarp-derived structures (Sukhorukov *et al*. 2021), such as persistent bracts in *Bougainvillea* or winged anthocarps in *Phaeoptilum*, which restrict its dispersal. Thus, the low records of *Belemia* species are likely due to life-history traits that reduce dispersal, leading to their current narrow distribution.

*Belemia fucsioides* has been included in the List of Endangered Fauna and Flora – Espírito Santo (Fraga *et al*. 2019) (https://iema.es.gov.br/especies-ameacadas/ameacadas). Since that report, botanical research has yielded only five observations of this plant at two known locations, and our investigation confirms the ongoing threats to its ecosystem. Some of the locations where *B. fucsioides* occur in Espírito Santo have been converted to sugarcane and *Eucalyptus* L’Hér. plantations. In fact, we did not find the species at the 2008 collection site on three consecutive attempts in 2018, 2021, and 2026 because the cultivated area had expanded into the forest fragment where it was previously collected. *Belemia cordata* faces similar conditions in the Cerrado in Tocantins state. *Eucalyptus* plantations and cattle activity pose a serious threat to the habitat of this species, which has been found only twice, growing either along pasture fences or on top of disturbed vegetation within these disturbed areas. Thus, in addition to threats from habitat loss and fragmentation, *Belemia* species also face competition from invasive exotic species, further worsening their survival challenges. Species from the grass family (Poaceae) and legume family (Fabaceae) were particularly commonly associated with *Belemia* species, including *Acacia mangium* and *Leucaena leucocephala* (Lam.) de Wit in the range of *B. fucsioides* in northern Espirito Santo, and *Mimosa* sp. in the range of *B. cordata* (Fig. 5D).

To better understand the status of *Belemia* and reduce threats to its species, we recommend monitoring habitat fragmentation and conducting studies to identify key biological factors that could prevent its establishment in natural areas, including investigations of its pollination biology, phenology, and ecology. It is also important to raise awareness of their endangered status among the local population and stakeholders involved in the economic activities that threaten the species. The fact that three or four of the records of *B. fucsioides* were found in protected areas (i.e., Reserva Biológica de Córrego do Veado) emphasizes the importance of these initiatives for species conservation. However, most other areas where the species has been collected are outside any protected area. Importantly, there is no conservation unit within the range of *B. cordata*, yet agricultural and forestry activities continue to increase in the region. These threats have endangered the beautiful *Belemia* plants—along with many other species. We urge policymakers, land managers, and local stakeholders to support botanical research, habitat monitoring, and conservation actions aimed at protecting these narrowly endemic species and their threatened ecosystems.

## Acknowledgements

I.L.C.N is grateful to Gabriel F. Rezende for helping with the permission to visit the Reserva Biológica Córrego do Veado (SISBIO license 59454) and Jovano F. Oliveira for helping during field collection. We also thank Maria F. Colodete and Adriana C. S. Cavalcanti (2008), Luiza F. Souza (2026), and Henrique S. Pereira (2026) for their support during field collections.

## Authors’ Contributions

I.L.C.N. and C.F.C.S.: conceptualization, field collection, data curation, formal analysis, funding acquisition, writing– original draft; E.F.S.R.: field collection, data curation, formal analysis, writing– original draft; D.V.G.: data curation, formal analysis, writing – review & editing; M.G.C.N.: field collection, locality data, historical information on the discovery of *Belemia cordata*, review & editing.; G.M.A., A.O.S., V.R.C.R.: field collection, writing – review & editing; V.A.: funding acquisition, writing – review & editing. All authors read and approved the final version of the manuscript.

## Data Availability

All data are contained in the manuscript and supplemental materials.

## Supplementary Material

The following online material is available for this article:

**Supplementary Table S1.** List of plant material collected and investigated in the present study.

**Supplementary Table S2.** Morphological comparison between *Belemia fucsioides* and *B.cordata*.

---

**Supplementary Table S1.** List of *Belemia* specimens collected to date and investigated in the present study.

| Taxa | Locality | Year Collected | Voucher and Herbarium | Geographical analysis | Morphological analysis |
| --- | --- | --- | --- | --- | --- |
| *B. cordata* | Ananás-Angico, Tocantins | 2016 | Nogueira 544 | Yes | Yes |
| *B. cordata* | Ananás-Angico, Tocantins | 2026 | Silva 130 | Yes | Yes |
| *B. fucsioides* | Espírito Santo | 1953 | Duarte 3704 | Yes | Yes |
| *B. fucsioides* | Espírito Santo | 1965 | Lanna-Sobrinho 1003 | Yes | Yes |
| *B. fucsioides* | Espírito Santo | 1965 | Belem 1460 | Yes | No |
| *B. fucsioides* | Minas Gerais | 1968 | Belem 3796 | Yes | Yes |
| *B. fucsioides* | Pedro Canário, Espírito Santo | 2008 | Sá 4887 | Yes | Yes |
| *B. fucsioides* | Pedro Canário, Espírito Santo | 2008 | Sá 4888 | Yes | Yes |
| *B. fucsioides* | Pinheiros, Espírito Santo* | 2009 | Folli 6510 | Yes | Yes |
| *B. fucsioides* | Mucurici, Espírito Santo | 2021 | Stehmann 6568 | Yes | No |
| *B. fucsioides* | Pinheiros, Espírito Santo* | 2021 | Cunha-Neto 16 | Yes | Yes |
| *B. fucsioides* | Pedro Canário, Espírito Santo | 2021 | Cunha-Neto 17 | Yes | Yes |
| *B. fucsioides* | Pinheiros, Espírito Santo | 2022 | Nunes 101 | No | No |
| *B. fucsioides* | Pinheiros, Espírito Santo* | 2022 | Nunes 101 | No | No |
| *B. fucsioides* | Pedro Canário, Espírito Santo | 2026 | Antar 6212 | Yes | Yes |

\* Reserva Biológica de Córrego do Veado

**Supplementary Table S2.**
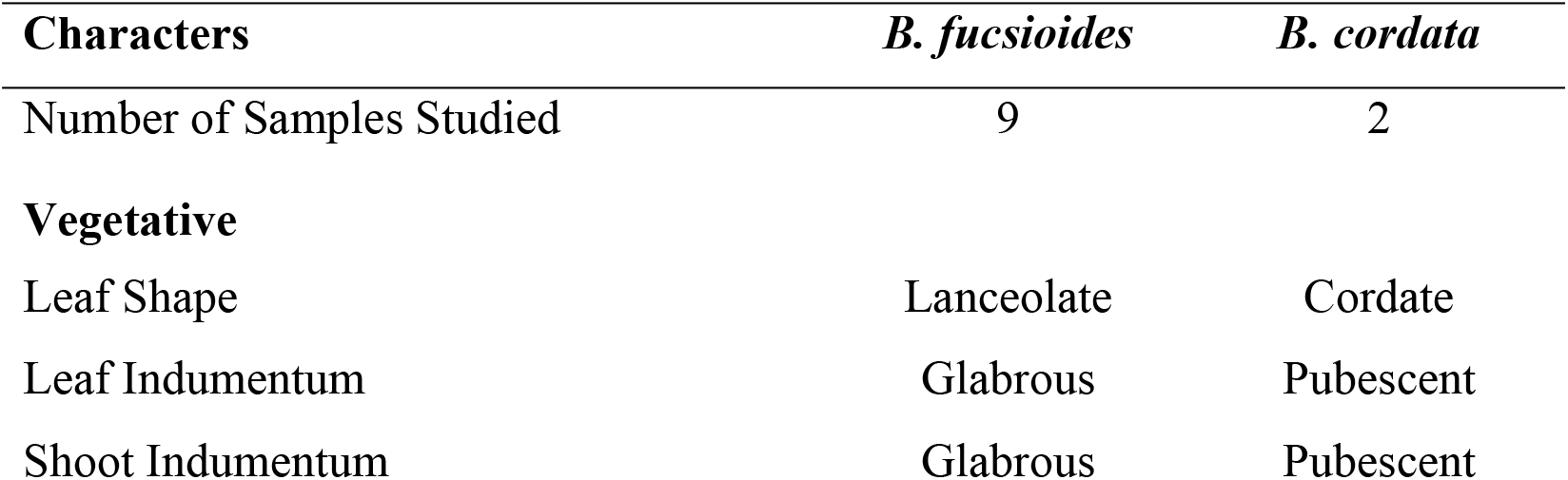
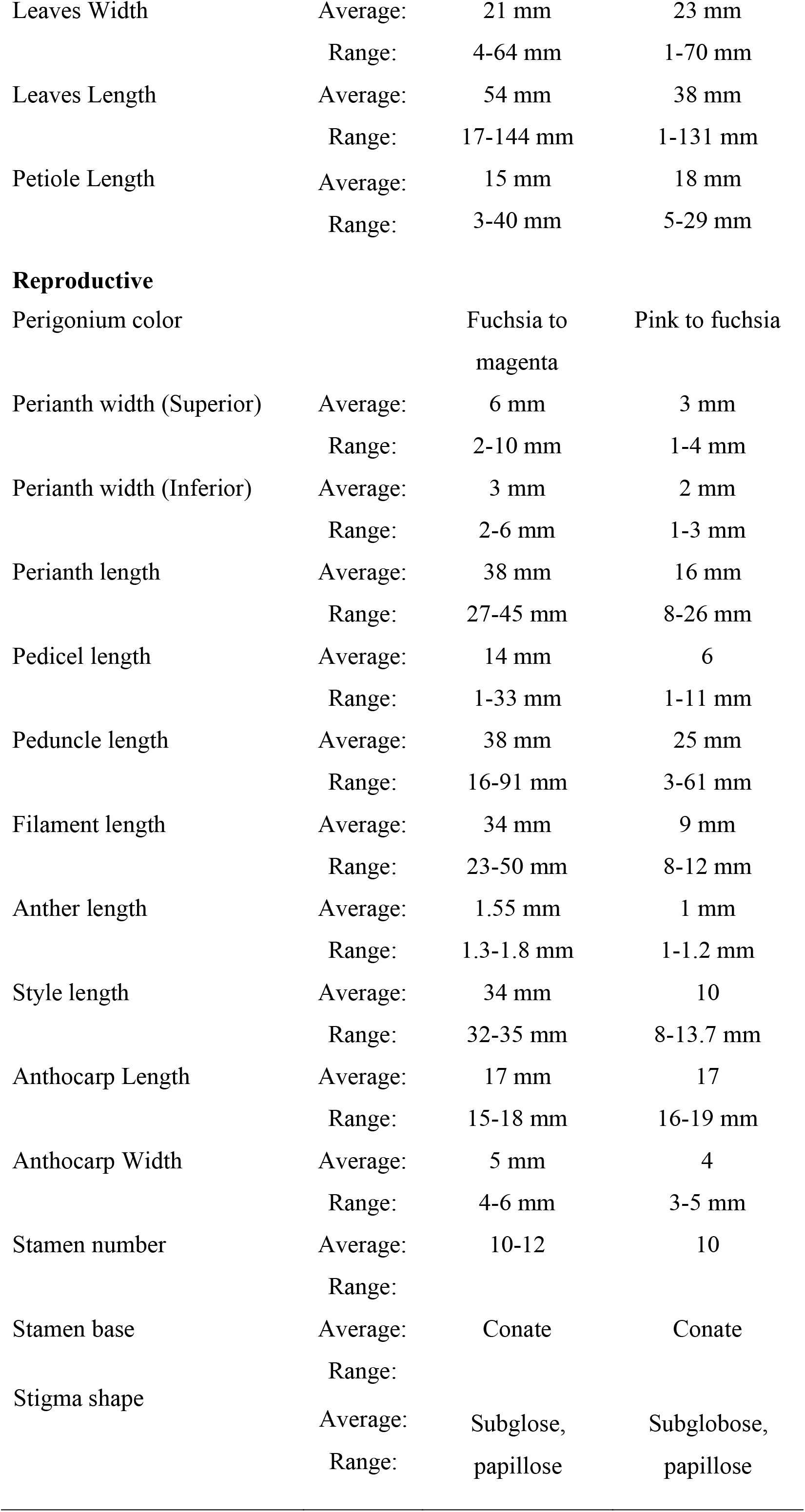
Morphological comparison between *Belemia fucsioides* and *B. cordata*.

| Characters | *B. fucsioides* | *B. cordata* |
| --- | --- | --- |
| Number of Samples Studied | 9 | 2 |
| **Vegetative** |  |  |
| Leaf Shape | Lanceolate | Cordate |
| Leaf Indumentum | Glabrous | Pubescent |
| Shoot Indumentum | Glabrous | Pubescent |

|  |  |  |  |
| --- | --- | --- | --- |
| Leaves Width | Average: | 21 mm | 23 mm |
|  | Range: | 4-64 mm | 1-70 mm |
| Leaves Length | Average: | 54 mm | 38 mm |
|  | Range: | 17-144 mm | 1-131 mm |
| Petiole Length | Average: | 15 mm | 18 mm |
|  | Range: | 3-40 mm | 5-29 mm |
| **Reproductive** |  |  |  |
| Perigonium color |  | Fuchsia to magenta | Pink to fuchsia |
| Perianth width (Superior) | Average: | 6 mm | 3 mm |
|  | Range: | 2-10 mm | 1-4 mm |
| Perianth width (Inferior) | Average: | 3 mm | 2 mm |
|  | Range: | 2-6 mm | 1-3 mm |
| Perianth length | Average: | 38 mm | 16 mm |
|  | Range: | 27-45 mm | 8-26 mm |
| Pedicel length | Average: | 14 mm | 6 |
|  | Range: | 1-33 mm | 1-11 mm |
| Peduncle length | Average: | 38 mm | 25 mm |
|  | Range: | 16-91 mm | 3-61 mm |
| Filament length | Average: | 34 mm | 9 mm |
|  | Range: | 23-50 mm | 8-12 mm |
| Anther length | Average: | 1.55 mm | 1 mm |
|  | Range: | 1.3-1.8 mm | 1-1.2 mm |
| Style length | Average: | 34 mm | 10 |
|  | Range: | 32-35 mm | 8-13.7 mm |
| Anthocarp Length | Average: | 17 mm | 17 |
|  | Range: | 15-18 mm | 16-19 mm |
| Anthocarp Width | Average: | 5 mm | 4 |
|  | Range: | 4-6 mm | 3-5 mm |
| Stamen number | Average: | 10-12 | 10 |
|  | Range: |  |  |
| Stamen base | Average: | Conate | Conate |
|  | Range: |  |  |

|  |  |  |  |
| --- | --- | --- | --- |
| Stigma shape | Average: | Subglose,<br>papillose | Subglobose,<br>papillose |
|  | Range: |  |  |

**Supplementary Figure S1.**
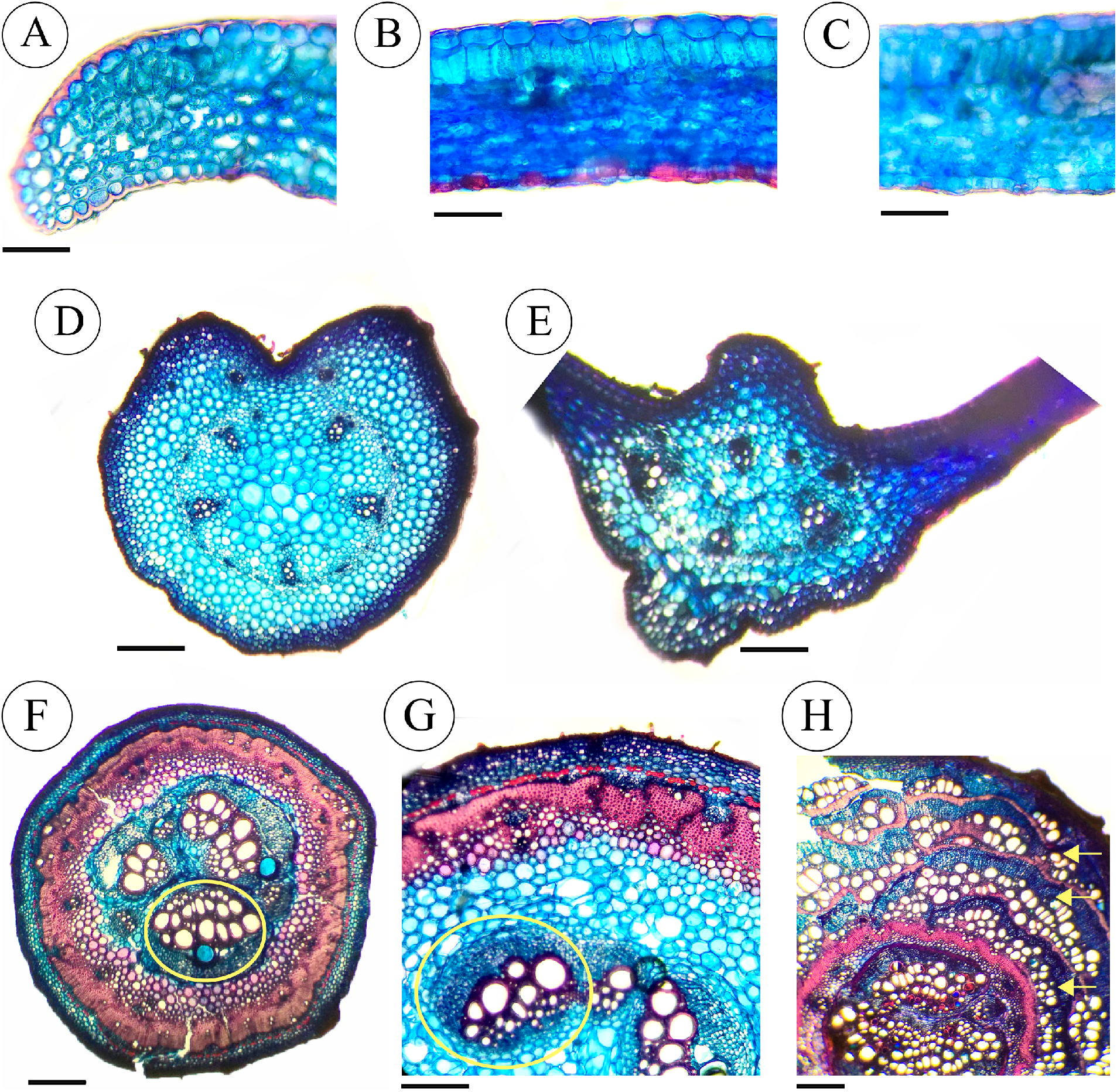
Vegetative anatomical features of *Belemia fucsioides* (*Cunha Neto 17*). All images are in cross-section. (A-C) Leaf blade. (A). Leaf margin, with visible cuticle. (B) Dorsiventral mesophyll. (C) Stomata on the abaxial surface. (D) Petiole; vascular system formed by five collateral bundles in the center, and two bundles towards the constriction zone. (E) Midrib; the vascular system includes 5-7 collateral bundles. (F-H) Stem. (F) Transition from primary to secondary growth; note the large “medullary bundles” (a type of vascular variant) located in the pith. (G) Detail of medullary bundles and the peripheral vascular cylinder with atypical cambium initiation. (H). Mature stem with “interxylary phloem” (a type of vascular variant), which forms bands of secondary xylem alternating with bands of secondary phloem (yellow arrows) derived from the activity of a single cambium with atypical activity. Circles (yellow) = medullary bundles. (A-F) 250 μm; (G) 100 μm; (H) 700 μm.

